# Position of *de novo* purine biosynthesis gene disruptions shapes purine-starvation phenotypes in *Saccharomyces cerevisiae*

**DOI:** 10.64898/2026.03.04.709599

**Authors:** Z. Ozolina, A. Kokina, A. Zile, E.T. Auzins, K. Pleiko, A. Kristjuhan, J. Liepins

## Abstract

Purine starvation induces a quiescence-like and stress-resistant physiological state in *Saccharomyces cerevisiae*, but it remains unclear whether this response is uniform across all disruptions of the *de novo* adenine biosynthetic pathway. Here, we used a pathway-position series of adenine-biosynthesis deletion mutants to test how the location of the metabolic blockage shapes cellular adaptation to purine starvation, using nitrogen starvation as a reference condition. Across mutants, purine starvation induced a shared core response marked by reduced growth, repression of ribosome-biogenesis and rRNA-processing modules, and accumulation of cells in G1/G0 phase. However, the magnitude and composition of the response varied systematically with pathway position. Early-pathway mutants showed the strongest desiccation tolerance and stress-associated transcriptional remodelling, whereas late-pathway mutants showed weaker protection. The *ade16Δade17Δ* double mutant was a distinct outlier, with altered carbon-flux distribution and limited stress resilience, suggesting that late-pathway disruption produces a physiologically different starvation state. Transcriptome analysis showed partial overlap with nitrogen starvation, but also pathway-position-dependent regulation of metabolic, ribosome-related, mitochondrial, and stress-response modules. Together, these results indicate that purine starvation is not a single physiological condition, but a set of related starvation states shaped by the position of the metabolic block within the purine biosynthetic pathway.

## Introduction

In rapidly changing environments, organisms such as yeast monitor external nutrient sources and adjust their metabolism accordingly. Typically, when the supply of essential compounds such as carbon, sulfur, phosphate, or nitrogen is depleted and starvation persists, a dormancy or “quiescence” program is activated. Under these conditions, the cell cycle is arrested in the G1/G0 phase, stress resistance mechanisms are induced, and cells can survive in this state for an extended period. Upon replenishment of the missing nutrients, cells can immediately resume proliferation (Sun and Gresham, 2021). However, starvation caused by the absence of a required metabolite does not always trigger the same regulated entry into a stress-resilient state. The effects of natural nutrient starvation differ significantly from those caused by auxotrophic starvation (Saldanha et al., 2004). During auxotrophic starvation, cells often fail to arrest their metabolism properly, leading to rapid cell death, as demonstrated in various studies. For instance, Boer et al. (2008) found that leucine- and uracil-starved cells undergo exponential death. Investigating lysine auxotrophy, Green et al. (2020) identified “nutrient-growth dysregulation” as a cause of rapid cell death during starvation. However, not all forms of auxotrophic starvation result in the same outcomes. By examining several cases, Lewis et al. (2024) and Petti et al. (2011) showed that phenotypes and survival rates vary greatly among different starvations. Thus, auxotrophic starvation cannot be treated as a single physiological phenomenon: both the missing metabolite and the architecture of the affected pathway may influence whether cells enter a stress-resistant cell state or undergo dysregulated reactions leading to death.

Purine starvation in *Saccharomyces cerevisiae* appears to represent an important exception to this pattern. Our previous studies showed that *ade8Δ* mutants, after a shift to purine-free medium, rapidly acquired a quiescent-like phenotype, already within four hours of starvation (Kokina et al., 2021). Kokina and colleagues also observed in red-pigment-forming mutants that purine starvation coincides with a dramatic increase in desiccation tolerance and a decrease in the budding index (Kokina et al., 2014). The purine synthesis pathway is highly conserved across all domains of life, particularly in eukaryotes (Chua and Fraser, 2020; Agmon et al., 2020), and purine moieties are central to cellular metabolism, signalling, and nucleic acid synthesis. Moreover, the inability to synthesise purines *de novo* is widespread in nature and is characteristic of many parasites, including *Leishmania spp., Trypanosoma brucei, Toxoplasma gondii, Plasmodium falciparum*, and microsporidia (Li et al., 2015; Boitz et al., 2012; Donaldson et al., 2014; Dean et al., 2016; Riegelhaupt et al., 2010). Thus, budding yeast provides a potential model for examining how eukaryotic cells respond to purine limitation, with possible relevance to purine-dependent parasites. However, most current studies of purine starvation have focused on individual auxotrophic mutants, leaving unresolved whether the observed physiological state reflects a general property of purine limitation or depends on the specific step at which the pathway is interrupted.

In budding yeast, the *de novo* purine synthesis branch consists of a series of 12 reactions from phosphoribosyl pyrophosphate to inosine monophosphate, catalysed by enzymes encoded by *ADE4* through *ADE16* and *ADE17* (Rebora et al., 2005), and then branches towards AMP and GMP synthesis, also connecting to the salvage pathway (Figure 1). This linear organisation makes the pathway a useful system for testing whether different positions of metabolic interruption produce equivalent or distinct starvation states. Deletion of any gene in this chain results in adenine auxotrophy, with one exception: both *ADE16* and *ADE17*, which catalyse the final steps in the linear part of the pathway, must be deleted simultaneously; in this case, the strain also becomes auxotrophic for histidine because of the metabolic link between the two pathways (Rebora et al., 2005). Two mutations in the purine biosynthesis pathway, *ade2Δ* and *ade1Δ*, have previously received particular attention and are widely used in laboratories (Bharathi et al., 2016; Stovicek et al., 2015). When the purine source is depleted*, ade2Δ* or *ade1Δ* cells accumulate AICAR, which in the presence of oxygen becomes intensely red, making these strains attractive for selection. Still, it remains unclear whether all adenine biosynthesis mutants produce the same quiescent-like phenotype under purine starvation. This raises the possibility that metabolic pathway architecture itself may determine the nature of the starvation response, such that different blocks within the same biosynthetic pathway generate distinct physiological states.

**Figure 1.**
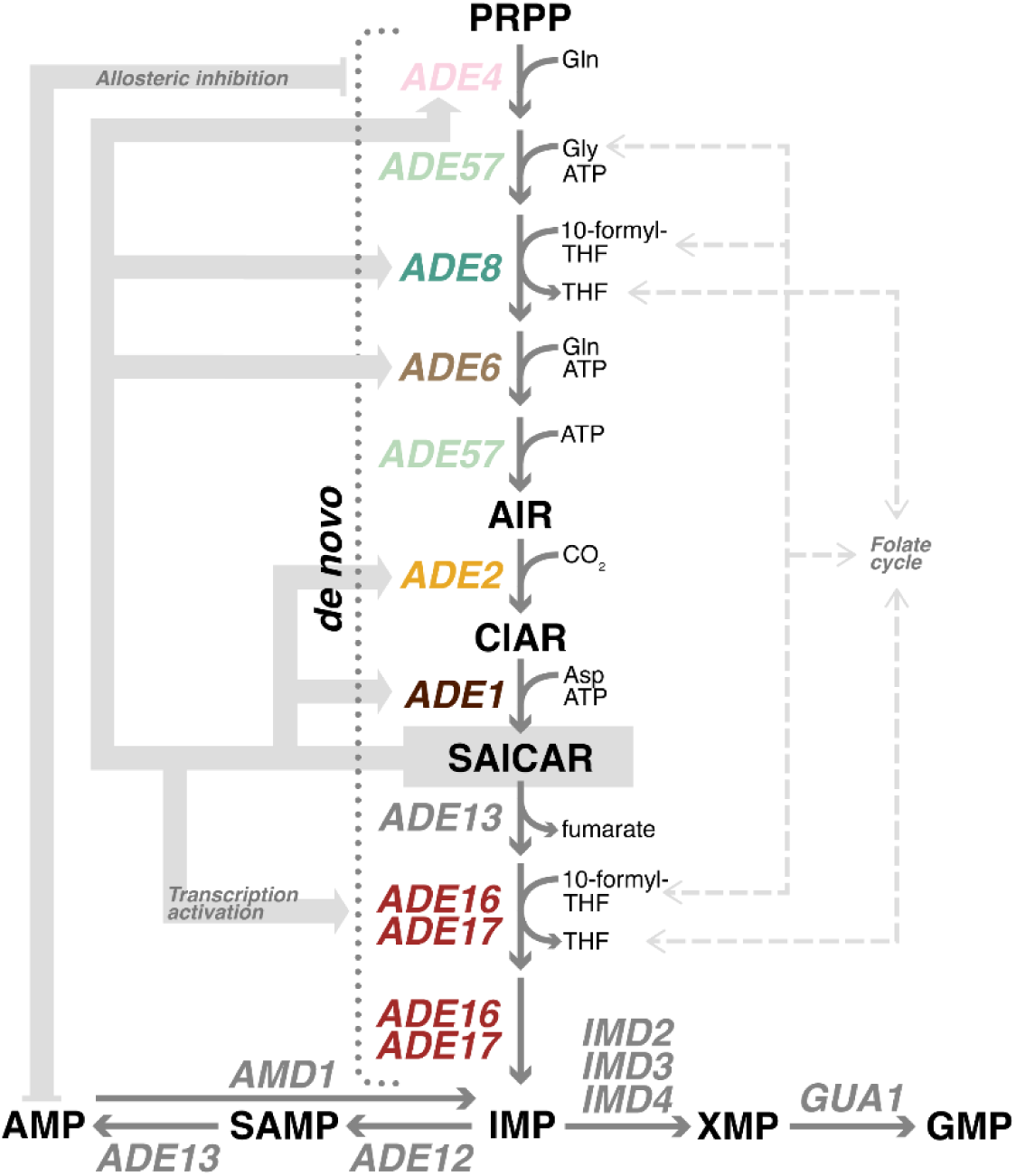
Yeast purine *de novo* biosynthesis pathway, adapted from Agmon et al. 2019.

We hypothesised that purine starvation in budding yeast engages a regulated starvation response that partly overlaps with nitrogen-starvation signalling, but that the final physiological state depends on where *de novo* purine synthesis is interrupted. To test this, we generated a series of adenine-biosynthesis mutants in a clean prototrophic CEN.PK background and compared their responses to short-term purine and nitrogen starvation. This experimental design allowed us to separate the general consequences of purine limitation from effects linked to the position of the metabolic block. Because auxotrophic markers can affect growth, physiology, and stress responses even in complete media (Mülleder et al., 2012; Kaplan et al., 2024), the use of prototrophic parental strains was important for reducing confounding effects. Nitrogen starvation was used as a reference condition because it represents a well-characterised natural starvation response and because adenine is a nitrogen-rich metabolite. By integrating growth, carbon-flux, stress-resistance, cell-cycle, trehalose, long-term survival, transcriptomic, and transcription-factor perturbation analyses, we tested whether interruption of a single conserved metabolic pathway generates one uniform starvation response or multiple pathway-position-dependent physiological states.

## Materials and methods

### Yeast strains

CEN.PK113-7A prototroph was used as the parental strain. Loss-of-function mutants affecting the purine biosynthesis pathway were generated using the method described by Janke et al. (2004). In brief, it involved the construction of a linear PCR fragment containing a kanMX4 cassette flanked by 50 bp homology regions of the target gene, followed by transformation through electroporation of the parental strain. Transformants were screened on YPD plates with G418 (200 μg/mL), and then the purine auxotrophy phenotype was confirmed on synthetic media with and without purine. Strain mating and tetrad dissection were used to obtain *ade16Δade17Δ*. Deletion of the target genes was confirmed by colony PCR (Looke et al., 2011) and by Sanger sequencing of the respective PCR fragment. All the strains used in this study are summarised in Appendix 1.

For CEN.PK prototroph and auxotrophic strains *ade4Δ*, *ade8Δ* and *ade2Δ,* additional mutations were introduced: transcription factor genes *GCN2, GCN4, GLN3, FHL1, MSN2, FKH1,* and *STE12* were replaced by the hygromycin B resistance gene. Transformants were screened on hygromycin B (200 μg/mL) plates, and strain identity was confirmed by Sanger sequencing.

Stock cultures were frozen at -75°C in 0.9% NaCl and 15% glycerol. Before use, they were maintained on agarized YPD plates at 4°C for up to five re-streaking times.

### Media and cultivation conditions

For all experiments, synthetic media (SD+) (Miller et al. 2013) with 2% glucose were used. For adenine auxotrophs, the media were supplemented with 100 mg L^-1 adenine. Further in the text, ade-refers to media without adenine, N- to media without ammonium sulfate. For the cultivation of *ade16Δade17Δ,* histidine (100 mg L^-1) was also added. Cells were cultivated at 30°C, 180 rpm, and the culture volume did not exceed one-fifth of the flask volume. Optical density was measured at 600 nm.

For obtaining growth curves, 96-well plates were used, and strains were cultivated in 200 µL of appropriate media in the multimode reader (Tecan M200) in at least three replicates. The cultivation cycle for the multimode reader was: 490-sec orbital (3.5 mm) shaking, waiting 60 s, and optical density measurement at 600 nm.

For all experiments, cells were grown in SD+ overnight culture first, then inoculated in an appropriate volume and grown overnight again to obtain exponential phase cells. Before media shifts, cells were washed with distilled water twice and then inoculated in fresh media (SD+, N- or ade-) with a cell density of approximately 1*10^7 cells/ mL and cultivated for the required time.

### Stress assays

After 4 h of incubation, cells were harvested by centrifugation, washed with distilled water twice, and aliquoted to 1 mL, OD_600_=2 (corresponds to 1*10^7 cells/ mL). Three aliquots were exposed to each stress. For thermal stress, cells were kept at 50°C for 7 min, for oxidative stress, cells were incubated in 10mM H_2_O_2_ for 20 minutes and afterwards washed twice with distilled water. For the desiccation tolerance test, cells were sedimented by centrifugation, the supernatant was removed, and pellets were air-dried in a desiccator at 30°C for 6 hours. After that, 1 mL of distilled water was added to resuspend the cells.

After appropriate stress treatments, cells were serially diluted, and dilutions were spotted on YPD plates to assess CFU/mL. To control cell losses during washing steps, the optical density of suspensions was measured, and CFU/ mL was adjusted for the OD value. YPD plates were placed in an incubator at 30°C, and after 24-48 h, CFU was counted.

For transcription factor mutants, another type of desiccation tolerance was adopted. Strains were grown as described before in SD+, N- and ade-media, and then cells were pipetted into two 96-well plates, three replicates for each strain in respective media. Optical density was measured in the multimode reader Tecan Infinite M200. After that, 150 μL were removed, and the plates were put in an Eppendorf vacuum concentrator, under an aqueous regime for 1 h. After that, plates without covers were put in an incubator at +30°C for 16 hours. 200 μL SD+ with adenine was added to each well, and plates were inserted for cultivation in the multimode reader (Tecan M200), and growth curves were obtained as described before.

### Flow cytometry

For flow cytometry, 1 mL samples were collected throughout cultivation (0 to 5th hour) in 1-hour intervals. For cells cultivated in SD+, only 0, 3rd and 5th hour samples were collected. Cells were suspended in ice-cold 70% ethanol, kept at 4°C overnight, the next day, centrifuged, and 50 mM citric acid was added. Then, the cells were centrifuged again and resuspended in citric acid with RNase A (10 µg/mL), sonicated 3×2 min, and left at 37°C overnight. After that, cells were centrifuged, and DRAQ5 (250 µM) was added. Cells were kept for at least 30 minutes at room temperature in a dark place. If needed, before analysis, cells were diluted in citric acid. Analyses were done by flow cytometry in an Amnis ImageStreamX MkII imaging flow cytometer, and dye was excited using a 642 nm laser. The target population was gated using the density plot based on the brightfield area and the brightfield aspect ratio. For each sample, data from at least 10 000 cells were collected. The cell cycle was determined from a histogram of linear Draq5 fluorescence intensity.

### Fermentation and metabolite analyses

Batch fermentations were performed in a Sartorius Q-plus system with a working volume of 0.3 L, a gas flow of 0.25 L min^-1, and an agitation rate of 400 rpm. Extracellular glucose, ethanol, acetate, and glycerol were measured simultaneously by an Agilent 1100 HPLC system with a Shodex Asahipak SH1011 column and quantified with a refractive index detector (RI detector RID G1362A). The flow rate of the mobile phase (0.01 N H_2_SO_4_) was 0.6 mL min−1; the sample injection volume was 5 μL.

Carbon dioxide evolution was recorded by an exhaust gas analyser (InforsHT) in parallel to the harvesting of metabolite samples. Glucose uptake, ethanol, acetate, glycerol, and biomass fluxes were calculated by the methodology used in Sauer et al. 1999.

Biomass concentration was determined as absorbance at 600 nm. Different conversion factors for OD_600_ to dry weight [g * L-1] were found depending on cultivation media: 0.1504 g*L-1*OD-1 for SD+, 0.1551 g*L-1*OD-1 for ade- and 0.1258 g*L-1*OD-1 for N-. Metabolite fluxes were calculated as mM *gDW-1 *h-1 in a similar way to Sauer et al. 1999.

### Trehalose measurements

Trehalose content was measured with an anthrone method similar to Trevelyan and Harrison (1956). Cell suspensions, collected at appropriate time points during fermentation, were centrifuged, supernatants were aspirated, and cell pellets were washed once with distilled water. 500 µL of 1 M trichloroacetic acid (TCA) solution was added to fix cells and extract trehalose. TCA-fixed cells were stored at -20℃.

To prepare extracts containing trehalose, TCA-fixed cells were centrifuged for 10 min at 12000 rpm, and supernatants were collected. Cell pellets were resuspended in 150 µL of distilled water, and suspensions were centrifuged for 10 min at 12,000 rpm. Supernatants were pooled.

A solution of 0.2% anthrone in 75% H_2_SO_4_ was used for the quantification of trehalose. Standard solutions with known concentrations of trehalose (serial dilutions of 0 - 8.0 g L^-1^ trehalose) were used. 10 µL of the recovered cell supernatant and 60 µL water in duplicate were pipetted into microtubes, and 330 µL of the anthrone solution was added. The microtubes containing the reaction mixtures were placed in a 100℃ heating block for 7 minutes, then cooled down to room temperature. 175 µL of the reacted mixture was pipetted into a 96-well plate in duplicates. Absorbance at λ=625 nm was measured. The trehalose concentration was calculated from the standard curve.

### Transcriptomics

After 4 hours of cultivation, 50 mL of cells were centrifuged and flash-frozen in liquid nitrogen and then stored at -75°C until RNA extraction. Extraction was done with the RiboPure™ RNA Purification Kit, yeast (ThermoFisher Scientific). For each strain and media condition, three biological replicates were prepared for sequencing. Raw data stored in the ENA database, in datasets PRJEB83590 (ade-), PRJEB83591 (N-) and PRJEB83589 (SD+).

RNA samples were prepared using a 3‘ mRNA-Seq Library Prep Kit (Lexogen) according to the manufacturer’s protocol. Libraries were sequenced on an Illumina MiSeq platform. Reads were quality filtered, trimmed, and processed as described below. Sequencing reads were quality filtered (Q=30), Illumina adapters and poly-A tails were removed and reads with a length of at least 100 nt were selected for further processing using cutadapt. Genes with less than 1 count per million (CPM) in fewer than 2 samples were filtered out; Benjamini and Hochberg’s method was used to calculate multiple comparison-adjusted p-values as false discovery rate (FDR). FDR < 0.001 and logFC > 1,5 were set as a threshold for significance.

Gene Ontology and KEGG analyses were done in RStudio 2024.04.2. The study was performed for the genes with adjusted p-value < 0.05 and the logFC difference of at least 1.5. The GO term analysis and KEGG analyses were done using the enrichGO function from the clusterProfiler package (Yu 2012, Yu 2024, Xu et al. 2024, Wu et al. 2021). As the assignment database, the *S.cerevisiae* genome from the package org. Sc. sgd.db was used (Carlson et al., 2014). GO enrichment analysis was done for all ontologies, using the Benjamini-Hochberg method for p-value adjustment. P- and Q-value cut-off was set to 0.05 and 0.2, respectively. The KEGG pathway analysis was done using the function enrichKEGG from the clusterProfiler package. Only those GO terms or KEGG pathways with a FoldEnrichment change of at least 1.5 were displayed.

### Statistics

All statistics and data analysis were done in RStudio 2024.04.2, unless stated otherwise. The normality of data was checked using the Shapiro-Wilk test. Statistically significant differences (p<0.05) were calculated using the Wilcoxon or t-test as indicated below each figure.

The half-life of long-term survival was calculated by fitting an exponential decay model to data points. To determine the empirical variance, bootstrapping was used. Data points were resampled 100 times, and the decay exponent was calculated. The decay exponent was then used to calculate the half-life- a time point at which half of the cell population remains alive. The mean and standard deviation were calculated from the bootstrap values.

## Results

### The growth dynamics of purine auxotrophic strains

We generated a set of adenine auxotrophic yeast strains in the prototrophic CEN.PK background to assess whether the position of the mutation in the adenine biosynthesis pathway influences strain characteristics. The only exceptional strain was *ade16Δade17Δ*, which required a double deletion because deletion of either adjacent gene alone does not induce auxotrophy. This strain is also a histidine auxotroph.

First, we tested the growth of all newly generated strains (Figure 2). They were grown overnight in SD+, then optical densities were measured and equalised, and cells were washed with distilled water. Then, strains were grown for 24 hours in 96-well plates to obtain growth curves. For growth in SD+, we tested two different concentrations of adenine - 100 mg L^-1 and 20 mg L^-1, the second one responds to a concentration that is widely used in other laboratories and suggested as sufficient for the growth of adenine auxotrophs (Agmon et al. 2020), but the largest one is five times higher. Also, growth without nitrogen and adenine was tested, as shown in Figure 2. In SD+, when adenine concentration is 100 mg L^-1, we can see that all strains except *ade16Δade17Δ* show very similar growth patterns. This strain has a significantly longer lag phase and grows more slowly than other mutants, even in SD+. The growth curve of this strain, when adenine concentration is 20 mg L^-1, is very similar. At 20 mg L⁻¹ adenine, growth differences became more apparent and broadly followed the position of the disrupted step in the pathway. *ade4Δ,* which has a mutation in the first step, shows the highest sensitivity, and the growth curve flattened earlier, leading to the lowest maximal optical density among the tested adenine auxotrophic strains (without including *ade16Δade17Δ).* Similar growth patterns as we see of the strain *ade2Δ* in media with adenine concentrations of 20 and 100 mg L^-1 are also shown in Yan et al. 2023. When nitrogen ((NH_4_)_2_SO_4_) was omitted, no strains showed significant growth, but with no adenine, auxotrophic strains also did not grow in this medium. In both situations, without needed nutrients, some strains showed a small increase in optical density. Most likely, these strains used internal reserves to sustain growth; however, it did not last long (Kokina et al. 2014).

**Figure 2.**
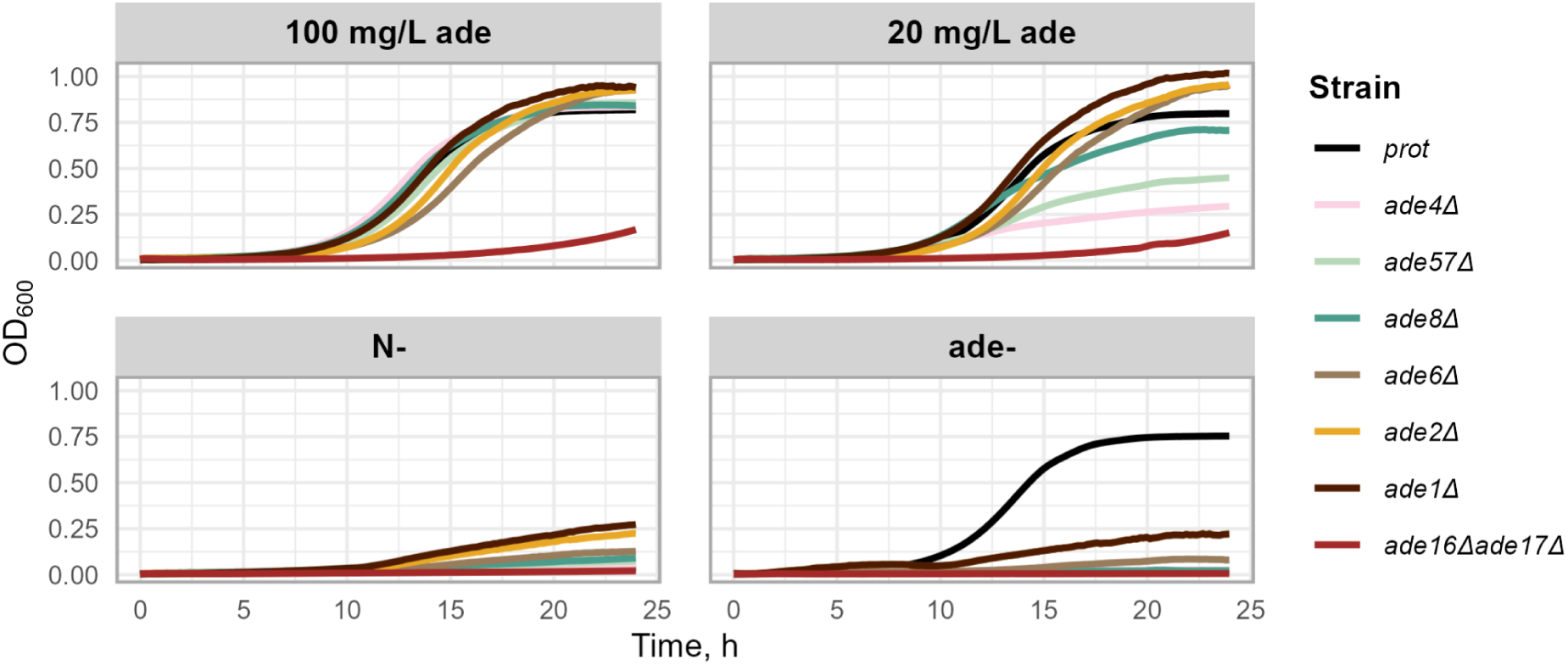
Growth curves of adenine auxotrophs used in this study in synthetic media with (100 mg L^-1 or 20 mg L^-1) and without adenine (ade-) or nitrogen source (N-). For *ade16Δade17Δ,* histidine (100 mg L^-1) was added. Strains with respective mutations here and also in the next figures are arranged in the order of the adenine synthesis pathway. Cultivations were carried out in 96-well plates; the graphs represent average results from 4 wells.

Interestingly, the strains that reached higher maximal optical density broadly followed the order of steps in the biosynthesis pathway, indicating that they may differ in their adenine requirements for optimal growth. To avoid differential effects of adenine limitation in subsequent experiments, we used 100 mg L^-1 adenine in SD+; for *ade16Δade17Δ*, histidine was added at the same concentration. Our results indicate that 20 mg L^-1 adenine is not sufficient to ensure unaffected growth of all mutants (Figure 2). Previously, some discussions have been raised on the optimal concentration of auxotrophic supplements (Gomes et al. 2007, Hanscho et al. 2012, Yan et al. 2023), and our results show that the amount of available purine modifies the growth and physiology of different purine auxotrophs.

### Carbon fluxes are redistributed during purine starvation

We conducted lab-scale fermentations to examine how purine starvation affects growth and the distribution of major carbon fluxes (Figure 3; see Materials and methods, Fermentation and metabolite analyses). Nitrogen starvation was used as a comparison condition. Growth results were similar to those obtained in 96-well plates. Figure 3A shows that growth rates in SD+ were similar across most strains; only the prototroph and *ade8Δ* grew slightly faster, whereas *ade16Δade17Δ* grew much more slowly, including under starvation conditions. However, the small growth observed in other strains under starvation conditions is likely due to a brief period of growth at the onset of starvation, similar to that seen in 96-well plates (Figure 2). Part of the increase in optical density may also reflect an increase in cell size during purine starvation, as reported previously (Kokina et al. 2021, Kokina et al. 2014).

**Figure 3.**
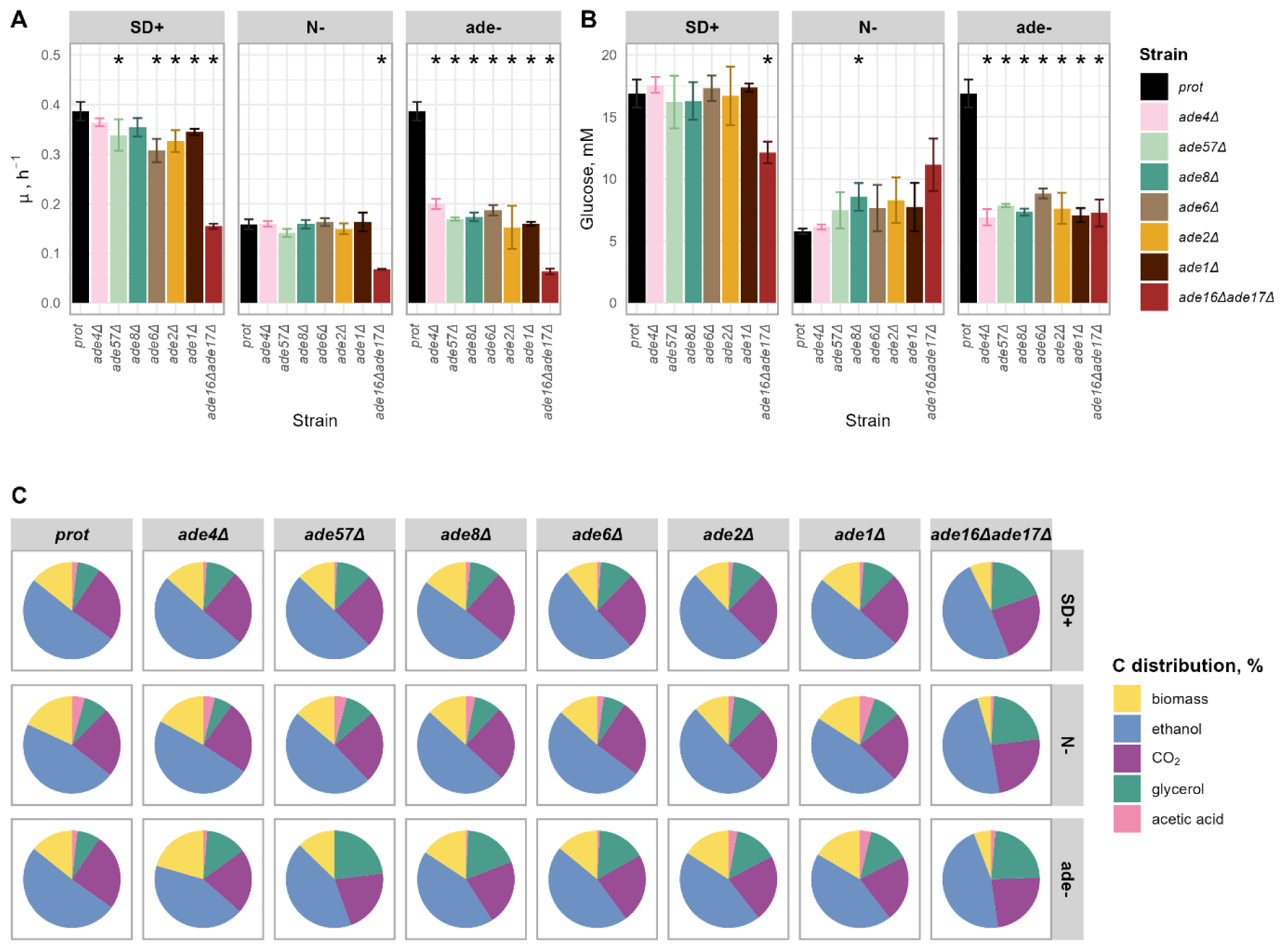
A Growth rate, h^-1^ of adenine auxotrophs in synthetic media with or without 100 mg L^-1 adenine and with or without nitrogen, strains cultivated for at least six hours in a Sartorius Q-plus fermentation system with a working volume of 0.3 L. B Glucose uptake, mM*h^-1^* gDW for the same strains, obtained during respective fermentation experiments. Stars denote statistically significant differences from prototroph (prot) in the respective condition (t-test, p<0.05). C Carbon flux distribution, showing the major carbon metabolites as % of metabolite flux in mCmol.

We measured the main carbon fluxes (Appendix 2) to get an overview of the changes in cell metabolism during starvation conditions. First, glucose uptake was calculated, and the results are summarised in Figure 3B, but the broader carbon-flux distribution is shown in Figure 3C. In SD+, the only strain with a significantly smaller glucose uptake rate was *ade16Δade17Δ* - in SD+, resembling N- glucose uptake rate for this strain, unlike the pattern seen in the other strains. As reported before, usually during starvation conditions, yeast tends to shut down its carbon metabolism and glucose uptake rates drop (Boer et al. 2008). This was also observed during purine starvation, but the glucose-wasting phenotype often observed during auxotrophic starvations (Boer et al. 2008, Petti et al. 2011) was not observed in purine-starved cells. In general, glucose consumption rates in ade- and N- are similar, approximately half as much as when cells are fast-growing. Thus, both purine and nitrogen starvation induced a broadly similar reduction in carbon uptake, consistent with a shared starvation-like metabolic program.

The distribution of consumed glucose was examined in more detail; all strains except *ade16Δade17Δ* showed similar patterns of carbon allocation to biomass. When starved, cells allocated only about one-third as much carbon to biomass formation as they did in full medium. This was true for both N- and ade- conditions, although carbon allocation to biomass was slightly higher in ade- than in N-. The biomass accumulation rate of the *ade16Δade1*7*Δ* strain in full media is similar to that of other strains in starvation conditions and even smaller in N- and ade- conditions. Also, the production of ethanol and, consequently, CO_2_, showed similar tendencies for all strains in respective conditions (around 22 - 29 mM / h*gDW in SD+, around 8-12 mM / h*gDW in ade- and N-). In this parameter, *ade16Δade17Δ* does not show significant differences. These results indicate that purine and nitrogen starvation similarly affect general carbon metabolism by slowing growth and reducing carbon investment in biomass formation and alcoholic fermentation.

Divergence among strains became most apparent when glycerol and acetic acid production rates were compared. Under nutrient-rich conditions, all adenine auxotrophs produced more glycerol (ranging from 3.19 mM/hgDW in *ade8Δ* to 5.45 mM/hgDW in *ade16Δade17Δ*) than the prototrophic strain (2.78 mM/hgDW). When starved, glycerol production generally decreased in all strains; however, auxotrophic strains still produced nearly twice as much glycerol when starved for adenine compared with the same strain starved for nitrogen. Notably, the double mutant *ade16Δade17Δ*, which carries mutations at the end of the adenine biosynthesis pathway, consistently showed the highest glycerol levels in all conditions (5.45 in SD+, 4.46 in N-, and 3.38 mM/hgDW in ade-). In glucose-grown cells, increased glycerol production is often associated with osmotic or redox stress responses (Petelenz-Kurdziel et al. 2013; Påhlman et al. 2001). In our data, however, glycerol elevation alone does not allow a specific interpretation, although it clearly distinguishes *ade16Δade17Δ* from the other mutants.

In contrast, acetic acid production showed the opposite trend. Across all auxotrophs, acetic acid levels were lower than in the prototroph under equivalent conditions. When starved for purines, strains with mutations early in the adenine pathway produced markedly less acetic acid (0-0.26 mM/hgDW) compared with nitrogen starvation (0.75-1.45 mM/hgDW). By comparison, *ade2Δ*, *ade1Δ*, and *ade16Δade17Δ* mutants produced similar amounts of acetic acid under both starvation conditions. The lower acetic acid production under purine starvation, especially in early-pathway mutants, further distinguishes ade- from N- and contributes to the pathway-position-dependent pattern. Thus, purine starvation produced a shared reduction in central carbon flux, but glycerol and acetate production revealed pathway-position-dependent metabolic states.

### Purine-starved cells are stress-tolerant and particularly desiccation-tolerant

It is already known that purine starvation causes higher stress tolerance in *ade2Δ* and *ade8Δ* auxotrophic yeast strains (Kokina et al. 2014, Kokina et al. 2021); similar effects are also seen in the intracellular parasite *Leishmania donovani* (Martin et al. 2016). We decided to check whether this happens for all mutants of the adenine biosynthesis pathway. Before stress, all strains were cultivated in SD+, N-, and ade- media for four hours, then washed, and the optical density was adjusted to identical values (OD_600_=2). Relatively short and severe stress treatments were used to assess how prepared cells are for environmental challenges after starvation. Results are summarised in Figure 4. As expected, the survival rate of exponentially growing cells cultivated in SD+ was low after stress, and this applies to all strains in this study, indicating that mutation in the adenine biosynthesis pathway alone does not alter the ability to withstand environmental stress when cells grow exponentially in full medium with an optimal amount of adenine.

**Figure 4.**
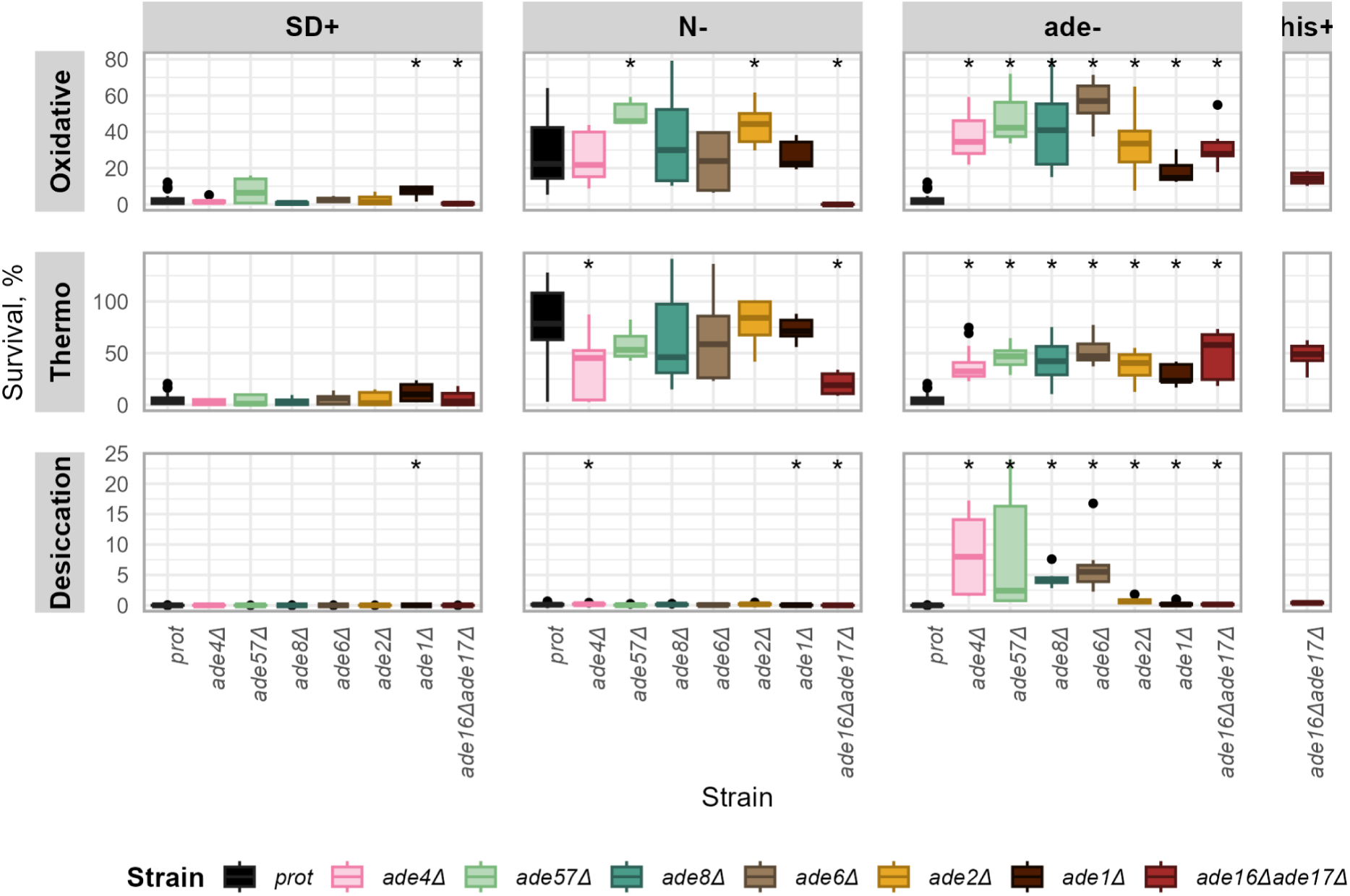
Stress tolerance of adenine auxotrophic yeast strains, viability measured as CFU, plated on YPD after stress, counted and compared with cell volume before stress. For oxidative stress, cells were kept in 10mM H_2_O_2_ for 20 minutes and afterwards washed twice with distilled water; for thermo stress, cells were stressed at 50°C for 7 min; desiccation - cells were dried in a desiccator for 6 hours at 30°C. Results represented in the graphs are averages from at least three biological replicates that were exposed to stress and at least eight technical replicates, plated and counted colonies after stress for each of them. Stars denote statistically significant differences from prototroph (prot) in the respective condition (t-test, p<0.05).

Nitrogen starvation, as previously widely described (Klosinska et al. 2011, Broach 2012), effectively increases stress resistance- our results show that even such a short period of starvation as we used- four hours- can turn on resilience mechanisms. Viability after oxidative and thermal stresses for almost all adenine auxotrophs is very similar to that of prototrophic strain and is high, ranging from 23% to 79%. After purine starvation, all adenine auxotrophs showed higher viability than the parental strain, ranging from 18% to 56%. Overall, the response to purine starvation in oxidative and thermal stress assays was broadly similar to that observed under nitrogen starvation. No significant differences were observed between strains from *ade4Δ* to *ade1Δ*; all responded within a similar range, indicating that these stresses mainly reveal a shared starvation-associated protective response.

Desiccation revealed a markedly different pattern from thermal and oxidative stress and provided the clearest separation between purine-starved mutants. This is a much more complex stress for a cell - it includes thermal, oxidative and osmotic stress simultaneously (Calahan et al. 2011). Survival after desiccation was lower than after heat or oxidative stress in all strains. Under this stress, purine-starved cells were much better protected, and viability ranged from 8% (*ade4Δ*) to 0.33% (*ade1Δ).* Strains with mutations at the beginning of the adenine biosynthesis pathway (*ade4Δ, ade57Δ)* showed higher viability than those with mutations at the end of it- viability for *ade2Δ* and *ade1Δ* is under 1% and is more similar to results for nitrogen-starved cells. Thus, unlike thermal and oxidative stress, desiccation clearly separates early- and late- pathway mutants, indicating that the position of the metabolic block strongly affects the protected state reached during purine starvation.

Strains with mutations at the end of the adenine biosynthesis pathway tended to show weaker stress resilience, and *ade16Δade17Δ* differed even more. Our growth results indicated that this strain grows more slowly in SD+, and its carbon fluxes also differ from those of the other strains (Figure 2, Figure 3). In general, stress resistance in this strain was lower than in other adenine auxotrophs, and it responded particularly poorly to nitrogen starvation. After thermal stress, viability was 20.43%; after oxidative stress and desiccation, no cells survived. After being starved for adenine (with and without histidine), *ade16Δade17Δ* showed better capacity to withstand all tested stresses than after being cultivated in nitrogen-free media; viability in this case is similar to *ade2Δ* and *ade1Δ.* Thus, although the strain retained the ability to respond to purine starvation, its phenotype clearly indicates that additional factors distinguish it from the other adenine auxotrophs.

### Purine auxotrophic starvation evokes G1/G0 arrest but does not fully explain stress resistance

Several physiological markers are associated with stress resistance and long-term survival. Here, we focused on two of them: cell-cycle arrest in G1/G0 and trehalose accumulation after four hours of cultivation. G1/G0 arrest is associated with entry into quiescence and is typically induced by starvation (Sun and Gresham 2021; Klosinska et al. 2011). Trehalose accumulation has also been linked to tolerance of several environmental stresses, although not all of them (Mahmud et al. 2010).

We asked whether the stronger stress-resistant phenotypes observed after four hours of starvation were followed by higher proportions of cells arrested in G1/G0 and by increased trehalose accumulation. They were not: both markers increased during starvation, but neither followed the pathway-position-dependent desiccation-tolerance pattern. Both purine and nitrogen starvation induced these classical starvation-associated markers; however, their magnitudes did not fully correspond to the differences in stress resilience observed between strains. A higher proportion of cells is arrested in G1/G0 during nitrogen starvation than during purine starvation, although stress resistance does not always follow this pattern. After four hours of nitrogen starvation, more than 70% of the population is arrested for all strains. During purine starvation, arrest does not happen in a fast and consistent way as in a medium without a nitrogen source. Still, after four hours, all auxotrophic strains have 60-75% of their population in G1/G0, see Figure 5.

**Figure 5.**
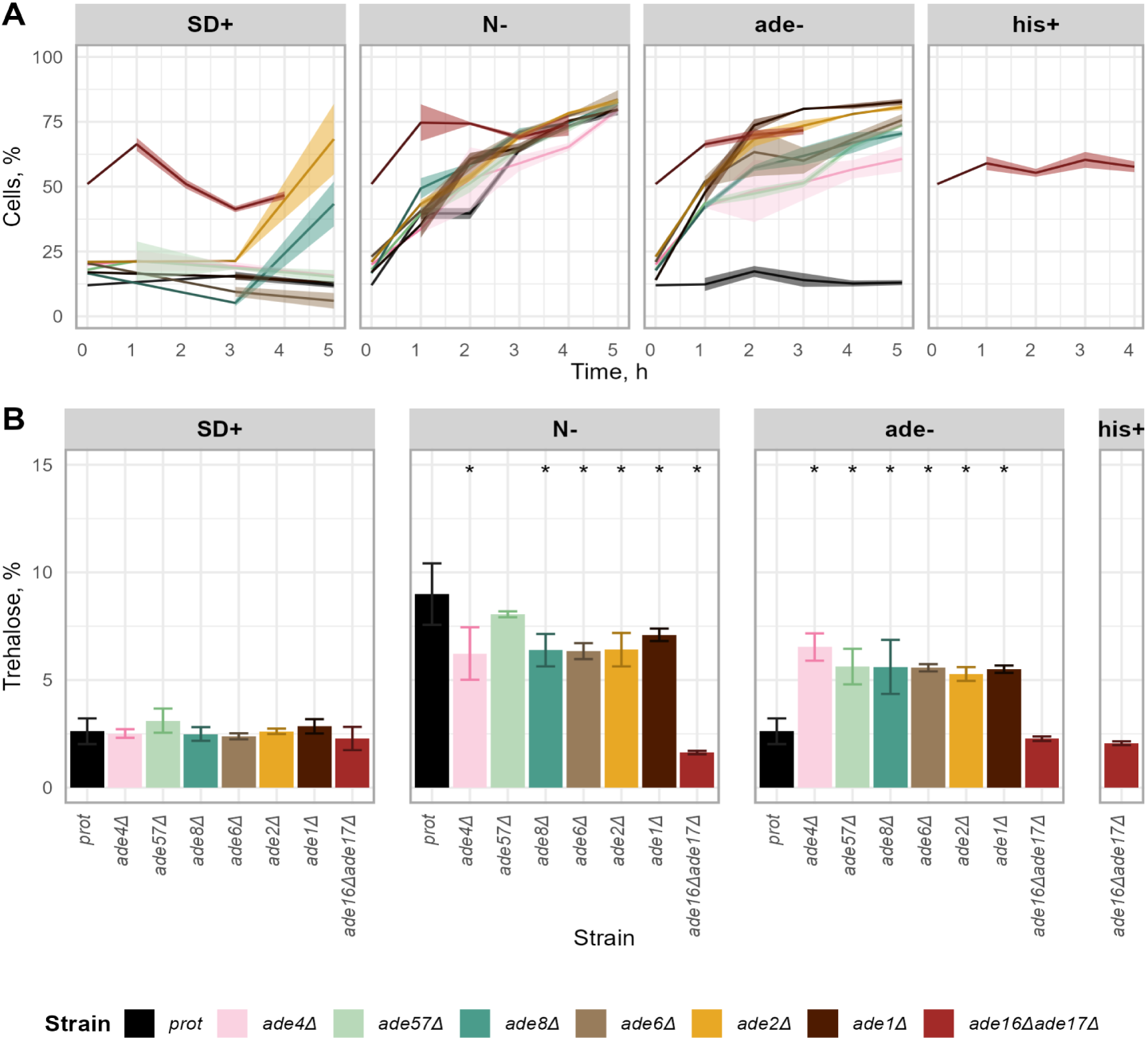
A Dynamics of cell cycle arrest in G1/G0 in adenine auxotroph strains during cultivation in SD+, N-, ade-. The lightly coloured area denotes the dispersion of the data points from three biological repeats. B Accumulation of trehalose after four hours of cultivation in SD+, N- and ade-media, % from dry weight. Error bars denote standard deviations from at least two biological and two technical replicates. Stars indicate statistically significant differences from prototroph (prot) in the respective condition (Wilcoxon test, p<0.05)

No significant differences appeared between strains during starvation conditions; notably, *ade4Δ*, which showed the strongest stress resistance under purine starvation (Figure 4), had the smallest G0/G1 arrested proportion of cells after four hours. Likewise, *ade16Δade17Δ*, which showed very weak stress resistance during nitrogen starvation, displayed a G1/G0 arrest pattern similar to that of the other auxotrophs. This strain and also *ade2Δ, ade8Δ* after the fifth hour of cultivation, had a fraction of the population also accumulated in G1/G0 even when conditions are favourable for exponential growth (SD+ media). Altogether, these results show that both nitrogen and purine starvation induce G1/G0 arrest, but the extent of arrest does not explain the differences in stress resistance, especially the strong divergence seen in desiccation tolerance (Figure 4).

Trehalose content was measured after four hours, the same cultivation time used before stresses. All strains cultivated in SD+ did not accumulate trehalose in significant amounts, accounting for 2-3% of biomass for all strains. Klosinska et al. 2011 indicated that nitrogen, but not carbon or phosphate starvation, induces trehalose accumulation. Our results are in agreement and confirmed that nitrogen-starved yeast cells do so- within four hours, they accumulated trehalose up to 6-7% of dry weight. Purine-starved cells accumulated similar amounts. Noticeably, the prototrophic strain during nitrogen starvation accumulated the highest amount of trehalose per gram DW. This amount was significantly higher than for all strains with mutations down from *ADE8*. Strain *ade16Δade17Δ* again showed significant differences from other adenine auxotrophs- an accumulated percentage of trehalose during nitrogen and purine starvations was small, similar to the results that other strains got after cultivation in SD+. This is consistent with the weak stress resistance of *ade16Δade17Δ* (Figure 4), but also reinforces the conclusion that neither G1/G0 arrest nor trehalose accumulation alone explains the previously observed pattern of stress resistance.

### Purine starvation affects the long-term survival of adenine auxotrophs

We continued the next experiments with four adenine auxotrophs representing different parts of the biosynthesis pathway, because strains with mutations in neighbouring steps had previously shown similar phenotypes. We chose *ade4Δ, ade8Δ, ade2Δ*, and the phenotypically distinct double mutant *ade16Δade17Δ*. We first asked whether these auxotrophic strains maintain viability during prolonged cultivation, so we cultivated these strains in SD+, N- and ade- for approximately 350 hours and monitored the viability of the population. Long-term survival is an important feature of the starvation response and has been reviewed in previous studies (Lewis et al. 2024; Petti et al. 2011). In our study, viability was expressed as CFU mL^-1 normalised to OD_600_=1 and compared with the SD+ four-hour time point, at which cells should be in exponential growth and were taken as 100% viable. Half-life was also calculated from this data. Because this method produces relatively high variance, the results should be interpreted mainly at the level of overall trends.

In SD+, all tested strains showed broadly similar survival dynamics: after approximately two days, about half of the population had lost viability, and after four to five days, more than 90% were non-viable (Figure 6A). In both ade- and N- conditions, starvation extended survival relative to SD+, with N- generally maintaining a slightly higher viable fraction. *ade16Δade17Δ* declined more rapidly than the other auxotrophs, although the overall trend of prolonged survival under starvation remained. Half-life values were broadly similar across strains, except for *ade8Δ* and *ade16Δade17Δ* in ade-, where they were shorter and closer to those of the prototroph (Figure 6B). We next asked whether the short-term desiccation-resistant state was maintained during prolonged starvation and whether it paralleled general long-term viability. Results (Figure 6A, indicated by dark lines) show that desiccation tolerance increased transiently under both starvation conditions and then declined. In SD+, only low desiccation tolerance appeared at later time points, most likely as nutrients became depleted. Under nitrogen starvation, desiccation tolerance peaked after 24 - 48 h and then gradually decreased. Under purine starvation, the same general pattern was observed, but desiccation tolerance was markedly higher, especially in early-pathway mutants. *ade4Δ* and *ade8Δ* retained higher viability after desiccation for longer than *ade2Δ*, whereas *ade16Δade17Δ* showed the lowest post-desiccation viability. Thus, long-term desiccation tolerance again supports the view that purine starvation produces a stress-resistant state, but this state is transient and remains dependent on the position of the metabolic block. Together, these results show that starvation prolongs survival relative to full medium, but long-term viability and desiccation tolerance do not fully overlap and therefore reflect different aspects of the purine-starvation phenotype.

**Figure 6.**
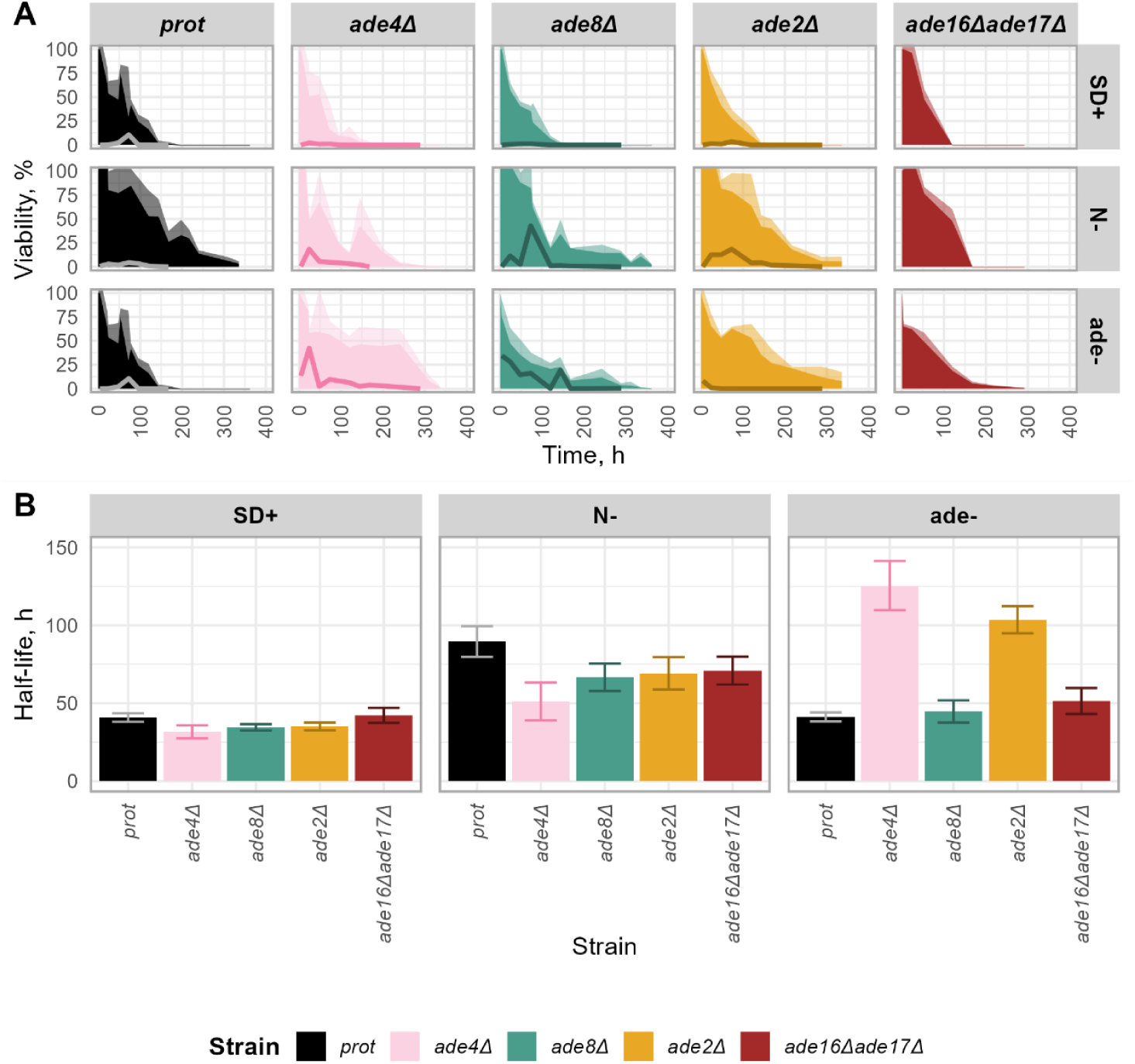
A Long-term survival in SD+, N- and ade- media, long-term viability (%) is shown as shaded areas; post-desiccation viability (%) is shown as dark lines. The lightly coloured area denotes the dispersion of the long-term survival data points from three biological repeats and at least eight technical repeats. B Half-life of long-term survival, h, calculated from cultivation in SD+, N- and ade-media. Error bars indicate the standard deviation of the data points after 100 bootstraps.

### Purine and nitrogen starvation share ribosome repression but trigger distinct metabolic and stress responses

To better understand the cellular response to purine starvation, we performed RNA-seq on cells cultivated for four hours in SD+, N- and ade-. Data were normalised against SD+, and genes with fold changes (FC) greater than 1.5 were analysed further. Volcano plots for all strains (Appendix 3) were first used to compare the overall scale and direction of transcriptional changes between strains and starvation conditions. A substantial part of the transcriptomic response overlapped between purine and nitrogen starvation, with ∼45 - 55% of significant genes shared between the two conditions. These shared genes were enriched in Gene Ontology categories related to stress response, carbohydrate reserve metabolism, and ribosome organisation (Figure 8). Thus, across all tested strains, the shift from complete medium (SD+) to adenine-free medium provoked a transcriptional programme that only partly overlapped with the canonical nitrogen-starvation response. Nitrogen starvation alone caused up-regulation of NCR-controlled permeases (*GAP1, MEP2, PUT4*) and autophagy markers (*ATG8, ATG14*), whereas these genes remained basal or modestly repressed in ade- conditions (Appendix 3), indicating a stronger NCR/TOR-associated signature in nitrogen starvation, as described previously (Beltran et al., 2004; Godard et al., 2007; Tesnière et al., 2015). Among the mutants, *ade4Δ*, which carries a deletion at the beginning of the adenine biosynthesis pathway, showed the largest set of differentially expressed genes (≈412), whereas *ade16Δade17Δ* showed the smallest response (≈173 transcripts) in the ade- versus SD+ comparison. Among the genes induced by purine starvation, *HSP12* was notable because of its established role as a hydrophilin associated with desiccation tolerance (Koshland and Tapia, 2019), consistent with the phenotype observed in Figure 4.

**Figure 7.**
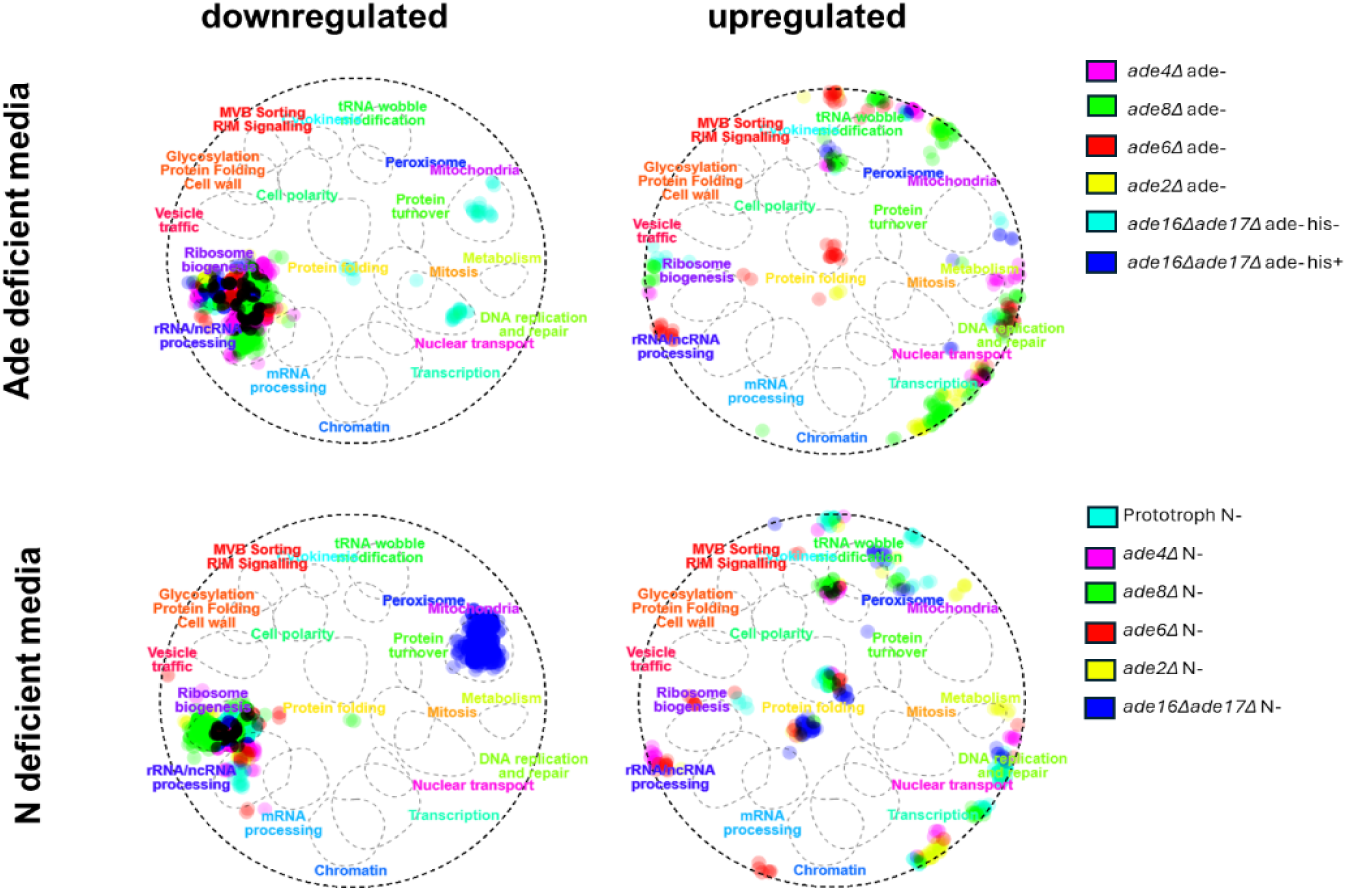
CellMap functional-interaction network, overlays of ≥1.5-fold differentially expressed genes in adenine auxotrophic strains and prototroph after 4 h adenine (ade-) or nitrogen (N-) starvation are depicted, showing up-regulated (left) and down-regulated (right) transcriptional modules for each condition.

**Figure 8.**
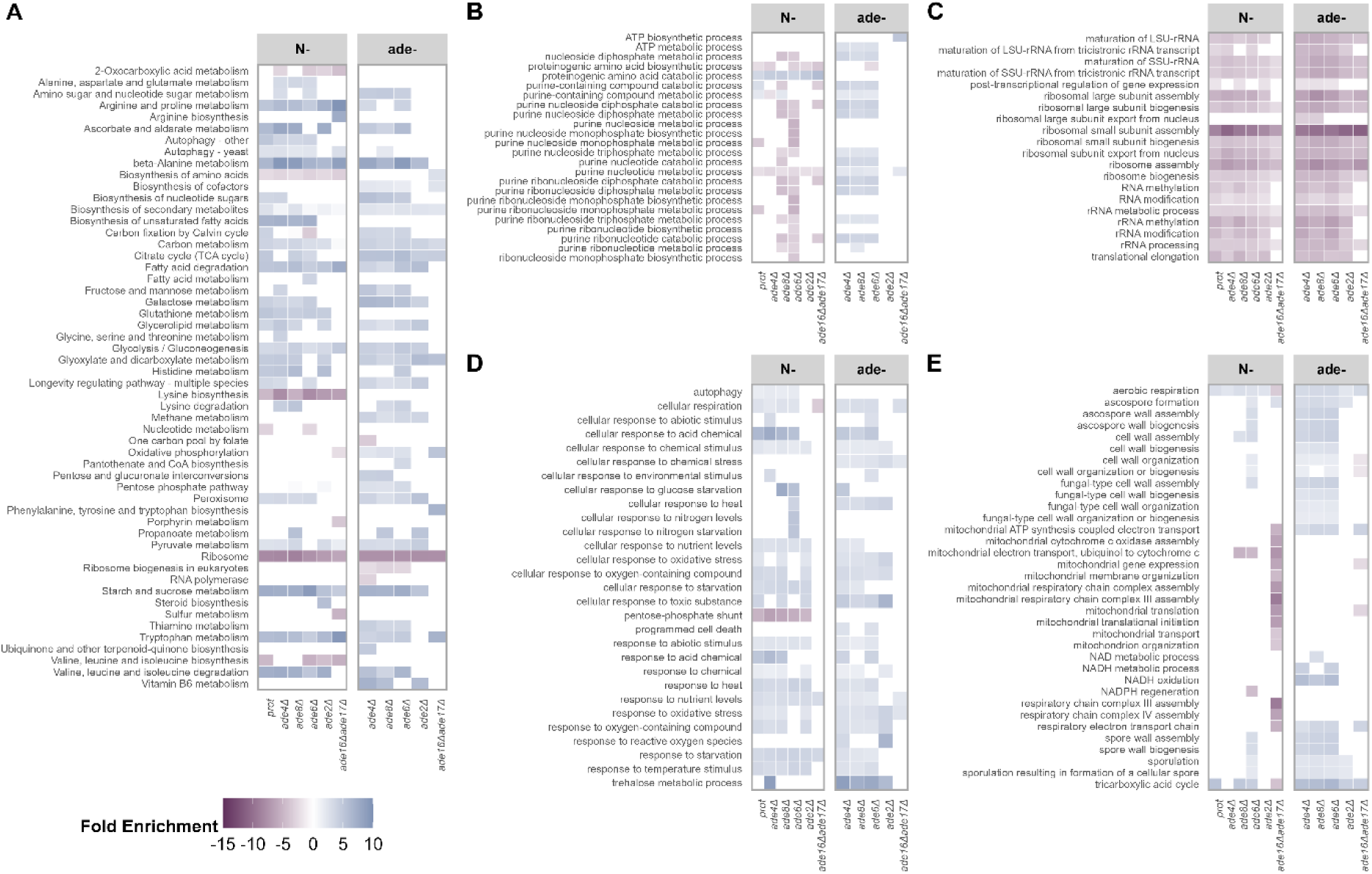
GO-Term Enrichment analysis of adenine biosynthesis pathway mutants, strains cultivated for four hours in SD+, N- and ade-, data normalised and fold change calculated against SD+, GO-terms arranged in five thematic blocks: A Amino-acid associated metabolism, B Purine metabolism associated processes, C Ribosome metabolism, D Stress associated processes, E Mitochondrial metabolism.

These patterns became clearer when differentially expressed genes were projected onto the CellMap functional-interaction network (Figure 7) (Costanzo et al., 2016). CellMap projection reinforced the main shared feature of both starvation conditions: strong downregulation of ribosome biogenesis and rRNA processing modules. Beyond this common core, *ade16Δade17Δ* showed an additional cluster of downregulated mitochondrion-peroxisome functions under nitrogen starvation, again distinguishing it from the other adenine biosynthesis mutants.

GO enrichment analysis recast the transcriptomic shifts into process-level patterns (Figure 8). Across the tested strains, purine starvation was associated with enrichment of central metabolic and stress-related processes, while both purine and nitrogen starvation strongly downregulated ribosome-related categories. Panel A summarises central and amino-acid-related metabolism GO terms and shows that adenine deprivation, irrespective of the mutant, enriches several central metabolic processes, including the pentose-phosphate pathway, galactose metabolism and TCA cycle. In the purine-metabolism-associated GO block, mild upregulation was seen in *ade4Δ, ade8Δ*, and *ade6Δ* during ade-, but disappeared in mutants at the end of the pathway, again indicating that the transcriptomic response depends on the position of the deletion. *ade16Δade17Δ* also showed strain-specific traits, including downregulation of sulfur metabolism and oxidative phosphorylation in N-. Panel C confirms that ribosome biogenesis and rRNA processing remain the most uniformly downregulated processes across both starvations, consistent with the canonical TOR-linked translation brake (Powers & Walter 1999). This panel again indicates that *ade16Δade17Δ* is an exception with clear differences in ribosome-related gene expression, showing much weaker downregulation of related GO terms. Stress-related GO terms were then examined (Figure 8D), and, as expected, most related processes were upregulated. Nevertheless, this panel also revealed a meaningful divergence: while both starvation conditions enriched responses to acid chemical, heat, oxidative stress, and starvation, only purine starvation showed a concerted rise in the trehalose metabolic process, despite similar trehalose accumulation under both conditions (Figure 5B). Only nitrogen starvation strongly downregulated the pentose-phosphate pathway, indicating that this pathway is relatively preserved during purine starvation. Because *ade16Δade17Δ* differed strongly in its transcriptional response, especially in genes related to mitochondrial metabolism (Figure 7), these categories were examined in more detail (Figure 8E). Panel E of Figure 8 shows that mitochondrial organisation and respiratory-chain assembly are broadly unperturbed in the remaining strains. Together, these data further distinguish *ade16Δade17Δ* from the other adenine biosynthesis mutants at the level of mitochondrial-related transcriptional response.

Together, the GO-term patterns support the transcriptomic hierarchy inferred from volcano plots (Appendix 3) and CellMap networks (Figure 7). Purine starvation preserved pentose-phosphate pathway capacity and was associated with a trehalose-related transcriptional response, two features that may contribute to the increased stress resistance of purine-starved auxotrophs. Across both starvation conditions, ribosome repression remained the strongest shared transcriptional response, indicating that translational shutdown may act as the backbone of the starvation program in budding yeast. This conclusion was further supported by transcription-factor analysis in YEASTRACT+ (Appendix 4). These analyses also indicated substantial overlap in the regulatory landscape of both starvation conditions, with Msn2/Msn4 among the major regulators of genes upregulated during purine starvation.

Based on the transcription factor (TF) analysis (Appendix 4), we generated a set of strains with deletions in candidate regulators of the purine-starvation response. The prototroph, *ade4Δ, ade8Δ*, and *ade2Δ* strains were used as parental backgrounds, and the genes encoding Fhl1, Fkh1, Gcn2, Gcn4, Gln3, Msn2, and Ste12 were deleted individually. Figure 9 quantifies how the removal of these transcription factors modulates the post-desiccation recovery of adenine-biosynthesis mutants. Post-desiccation recovery was quantified as the time required to reach OD_600_=0.3 and displayed as a heatmap (Figure 9). Panel A confirms the baseline pattern: after desiccation, all strains starved for purines resume growth more rapidly (≤ 8 h to reach OD_600_= 0.3; pale tiles) than those pre-starved for nitrogen, while strains grown in full media (SD+) show the longest lags (> 20 h; dark tiles). Panel B shows that deletion of any single regulator does not uniformly abolish this cross-protection, but instead creates genotype- and condition-specific delays. Under nitrogen starvation, some deletions produced longer recovery delays, whereas under purine starvation, the advantage of most mutants was retained. Loss of *MSN2* had little effect, consistent with previous reports that Msn2/4 are not always essential for desiccation tolerance under specific metabolic states (Calahan et al. 2011). None of the tested transcription-factor deletions had a strong and uniform effect across all strain backgrounds. Thus, the desiccation-resistant state induced by purine starvation appears to depend on a robust, multifactorial regulatory program that only partly overlaps with nitrogen-starvation signalling and canonical STRE-gene activation.

**Figure 9.**
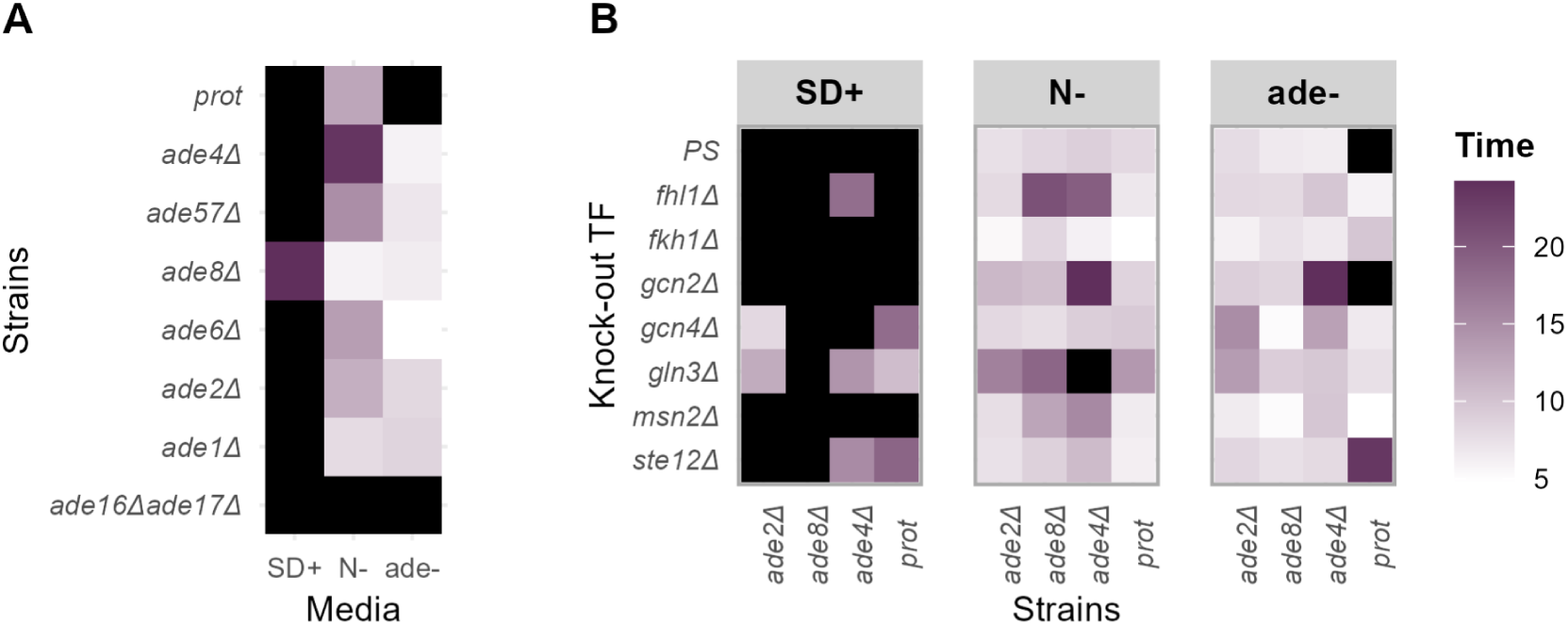
The time needed to reach OD_600_=0.3 after desiccation stress for adenine biosynthesis pathway mutants with deleted genes coding selected transcription factors: A parental strains, B strains with disrupted genes of selected transcription factors, black colour indicates time longer than 24 h.

## Discussion

In this study, we show that the outcome of purine starvation depends not only on the interruption of the *de novo* purine biosynthetic pathway but also on where it occurs. The response, therefore, appears not only to be a general reaction to growth limitation but also to include an organised starvation programme. In *S. cerevisiae*, the position of the disrupted step modulates stress tolerance, cell-cycle arrest, carbon-flux distribution, and transcriptional state. Since purine auxotrophy is widespread in nature, these findings may have broader relevance for understanding how cells respond to the limitation of essential metabolites.

### Purine starvation resembles a regulated natural starvation

Recognised nutrient stresses, including nitrogen and carbon starvation, activate signalling pathways that slow growth and increase stress resistance in yeast. Our data suggest that purine starvation is not simply a dysregulated auxotrophic defect, but a condition that can trigger features of regulated starvation response. Across strains we tested, the clearest shared transcriptomic signature was a repression of functions related to biosynthetic investment, especially ribosome biogenesis and rRNA processing, together with induction of stress-response genes (Figure 8C). From this viewpoint, purine starvation resembles nitrogen starvation. This overlap is biologically plausible because purines are nitrogen-rich metabolites, so purine limitation may intersect with pathways that monitor nitrogen availability and biosynthetic capacity.

Purine-containing molecules, including ATP, ADP, AMP, GTP, GDP, GMP, NAD⁺, NADP⁺, FAD, and S-adenosyl-L-methionine, are central to cellular metabolism. A lack of purine supply, whether caused by interruption of *de novo* synthesis or depletion of exogenous purines, can therefore affect several essential processes simultaneously, including energy metabolism, DNA and RNA synthesis, and cofactor metabolism. Moreover, purine-containing molecules are also linked to nutrient-dependent signalling, including pathways that respond to carbon and nitrogen availability (De Virgilio & Loewith, 2006). Previous results showed an approximately fourfold reduction in adenine nucleotide pools during purine starvation in *ade8Δ* cells, yet these cells still acquired increased tolerance to complex stress conditions (Kokina et al., 2021), consistent with the broader stress-tolerance pattern observed here (Figure 4). Low cellular ATP can suppress sugar uptake, linking energy charge to glycolytic flux (Schuddemat et al., 1988). Our carbon-flux data are consistent with this link and distinguish purine starvation from leucine or uracil auxotrophy, where cells continue glucose consumption, show ethanol overflow or “glucose-wasting,” and fail to arrest the cell cycle promptly (Boer et al., 2008; Petti et al., 2011). By contrast, purine-starved cells reduced carbon flux and accumulated in G1/G0 within the first hours of starvation, approaching the arrest pattern observed during nitrogen starvation (Figures 3 and 5A). These features support the idea that purine depletion can be translated into an organised starvation-like state rather than only into metabolic failure.

A similar purine starvation - associated phenotype has also been reported in other eukaryotic adenine auxotrophs, including intracellular parasites that rely on purine acquisition from the host cell (De Koning et al., 2005). Most of these parasites have lost the *de novo* purine biosynthetic pathway entirely and therefore depend on purine salvage. For example, when starved for purines, *Leishmania donovani* reduces intracellular ATP consumption, increases long-term survival, and arrests in G1/G0 (Martin et al., 2016). Studies in *Leishmania donovani* and *Crithidia luciliae* suggest that purine - dependent parasites can adjust adaptive responses to purine limitation (Alleman and Gottlieb, 1996; Carter et al., 2010). These parallels suggest that purine limitation can be sensed as a regulated metabolic stress in different eukaryotic systems, even though the underlying signalling mechanisms may differ. Understanding this response in budding yeast may therefore help to understand key principles of purine-limitation biology that are relevant to purine-dependent organisms and, indirectly, to therapeutic targeting of purine metabolism. Taken together, these observations support the view that purine depletion is not merely a dysregulated auxotrophic defect, but can trigger a regulated natural starvation-like response with reduced carbon flux, cell-cycle arrest, and increased stress tolerance.

### Why the position of disruption in the purine pathway matters

The phenotypic effect of a gene deletion depends not only on the gene itself, but also on its pathway and regulatory context (Caudal et al., 2022; Turco et al., 2023). Since previous studies have often focused on individual mutations (Rébora et al., 2005; Petti et al., 2011; Kokina et al., 2021), we compared deletions across the adenine biosynthesis pathway to test whether the position of the metabolic block affects the starvation phenotype (Figure 1). Although all *adeΔ* strains used here are adenine auxotrophs, they did not enter an identical physiological state; this became clear only when mutants were compared across the pathway rather than studied one by one. This difference was already visible in growth curves at 20 mg L^-1 adenine, where sensitivity broadly followed the order of the disrupted pathway steps (Figures 1 and 2). After purine starvation, most mutants showed broadly similar resistance to oxidative and heat stress, whereas desiccation tolerance separated the mutants more clearly. This phenotype separated early- and late-pathway blocks: strains disrupted near the beginning of the pathway, such as *ade4Δ* and *ade57Δ*, survived desiccation better than late-pathway mutants such as *ade2Δ* and *ade1Δ*, while *ade16Δade17Δ* behaved as a distinct outlier (Figures 4 and 6). This pattern is consistent with the idea that disruptions at different positions generate different intracellular metabolites and feedback signals. Early-pathway blocks may allow a more direct shutdown of growth and translation because fewer downstream intermediates are produced. By contrast, late-pathway blocks may allow the accumulation of intermediates such as AIR, CAIR, or AICAR, which are known to have a feedback effect on the regulation of ADE genes (Rébora et al., 2005), potentially modifying the starvation programme. This position-dependent accumulation of pathway intermediates, including AICAR, which can influence Bas1p-dependent ADE-gene regulation (Rébora et al., 2005), may be one of the key principles determining the response to starvation. In *ade2Δ* and *ade1Δ* strains, accumulation of AIR and CAIR has been linked to toxicity, including through conjugation with glutathione, which could interfere with oxidative-stress resistance and contribute to weaker desiccation tolerance. Such reactions may increase glutathione demand and thereby affect redox-stress management during starvation (Bharathi et al. 2016). Consistent with pathway-position-dependent feedback, the transcriptomic data showed graded differences in several GO-term blocks. The purine metabolism GO pattern supports this interpretation. In ade- medium, the mild upregulation of purine-pathway processes observed in early and middle mutants disappeared toward late deletions and was weakest in *ade16Δade17Δ* (Figure 8B). This suggests that block position reshapes transcriptional feedback rather than simply removing nucleotide supply. It may therefore influence which secondary metabolites and regulatory signals are produced, and which parts of the stress-response and quiescence programmes are involved. Taken together, our data show that adenine auxotrophy is not a single physiological perturbation, but a set of related metabolic disruptions that can produce distinct starvation phenotypes. Therefore, interpretation of purine-starvation phenotypes, including comparisons with natural starvation responses or purine-dependent parasites, should specify where *de novo* synthesis is interrupted, not only whether it is interrupted.

### The phenotype of ade16Δade17Δ is slightly atypical

Among the mutants analysed here, *ade16Δade17Δ* behaved as a distinct outlier rather than simply as the final point in the pathway-position series. This genotype is also histidine auxotrophic due to the metabolic coupling between purine and histidine biosynthesis (Rébora et al., 2005), and this additional feature should be considered when interpreting its phenotype. Although histidine was supplied in the media, this additional auxotrophy means that *ade16Δade17Δ* should be interpreted with caution. The strain was already exceptional in SD+: while most *adeΔ* mutants grew similarly with 100 mg L⁻¹ adenine, *ade16Δade17Δ* grew more slowly and showed an extended lag phase (Figure 2). The same distinction was observed in fermentation experiments- *ade16Δade17Δ* was the only strain with reduced glucose uptake already in SD+, resembling the lower uptake seen in other strains, mainly during starvation (Figure 3). At the same time, it showed the highest glycerol production across all tested conditions, suggesting an additional osmotic or redox burden. Phenotypically, this baseline difference was accompanied by weaker acquisition of stress resilience after starvation. Most strikingly, cells of *ade16Δade17Δ* gained small stress resilience ability from nitrogen starvation, with no detectable survival after oxidative stress or desiccation, whereas their response in ade- was closer to that of late- pathway mutants (Figure 4). This phenotype cannot be explained simply by failure to arrest the cell cycle. Under starvation, *ade16Δade17Δ* halted their cell cycle at G1/G0 similarly to the other auxotrophs (Figure 5A). However, trehalose accumulation remained low in both N- and ade-conditions, close to SD+ levels, which is consistent with its weaker stress resistance (Figures 4 and 5B). The transcriptomic data provide further clues to why this strain behaves as an outlier. In CellMap projections, repression of ribosome biogenesis and rRNA processing, which formed the shared backbone of the starvation response in other strains, was weaker in *ade16Δade17Δ* (Figure 7). In addition, this strain showed strain-specific downregulation of mitochondrial, peroxisomal, and respiratory functions under nitrogen starvation, as evidenced by both CellMap and GO enrichment for oxidative phosphorylation and sulfur-related terms (Figures 7 and 8E). The purine metabolism GO pattern also supports this interpretation: the mild upregulation observed in early and middle mutants disappeared toward late deletions and was weakest in *ade16Δade17Δ* (Figure 8B). Altogether, *ade16Δade17Δ* appears to execute part of the purine starvation-like response, but its phenotype likely reflects the combined consequences of a late block before IMP, purine-histidine pathway coupling, altered physiology, and possible changes in AICAR- or folate-linked metabolism.

### Increased desiccation tolerance is a specific feature of purine starvation

Desiccation is a multifactorial stress that combines osmotic, thermal, and oxidative challenges. As water is removed, cytoplasmic crowding increases, ionic balance is disturbed, and proteins and membranes become vulnerable to oxidative and structural damage (Capece et al. 2016; Calahan et al. 2011). This complexity was also reflected in our data. After 4 h of starvation, both N- and ade- conditions increased resistance against heat and oxidative stress, but they differed strongly in survival after desiccation (Figure 4). Nitrogen-starved cells remained below 1% viability after desiccation, whereas purine-starved mutants reached substantially higher survival, with the strongest resilience in early-pathway mutants, with survival reaching approximately 8% in *ade4Δ* and falling to approximately 0.33% in *ade1Δ*. After 24 h of starvation, desiccation tolerance increased further before declining during prolonged cultivation. Our data indicate that purine depletion induces a stress-resistant state that is especially evident under desiccation, where survival requires tolerance to several stresses at once.

In the transcriptomic data, several genes upregulated in ade- conditions encode proteins previously linked to desiccation or stress survival, including *HSP12* and *HSP26* (Appendix 3). Hsp12p and trehalose can act together to support viability after severe desiccation, and loss of both strongly compromises survival (Kim et al., 2018). In our data, purine starvation induces trehalose accumulation and also upregulates *HSP12*, suggesting a plausible route to protein stabilisation during drying and rehydration. Hsp26p may provide an additional recovery mechanism after rehydration. Although this has not been directly tested for desiccation in our study, Hsp26p has been implicated in protection during ethanol and heat stress, supporting its possible contribution (Ungelenk et al., 2016).

However, desiccation tolerance is unlikely to be explained by a single effector. Our transcription-factor deletion panel also argues against a single-master-regulator explanation for the purine-starvation phenotype (Figure 9). This is consistent with Calahan et al. (2011), who showed that single deletions affecting trehalose synthesis, hydrophilins, osmotic and salt responses, ROS detoxification, or DNA repair can leave desiccation tolerance largely intact. Instead, a notable influence for acquiring strong tolerance was mentioned to rely on respiration and on a combined activity of Msn2 and Msn4. This fits the broader view that the Msn2/4-dependent environmental stress response is particularly important for cross-protection against later severe stress, but is not always required for basal tolerance to a single stress (Berry and Gasch, 2008). Our data are consistent with this model: deletion of individual regulators produced genotype- and condition-specific recovery delays, but most purine-starved strains retained their post-desiccation recovery advantage. Only a small subset, including *ade4Δgcn2Δ, ade4Δgcn4Δ*, and *ade2Δgcn4Δ*, showed clearly prolonged lag (Figure 9). The limited effect of *MSN2* deletion is also consistent with partial redundancy between Msn2 and Msn4, although this would need direct testing with double mutants.

Our observations fit broader desiccation literature showing that desiccation tolerance is inducible and linked to nutrient signalling, respiration, and translation state, while general stress response activators alone cannot fully account for resistance (Calahan et al., 2011; Welch et al., 2013). The purine-starvation advantage becomes clearer when viewed not as the action of one effector, but as a system-level state combining protection before drying with recovery capacity after rehydration. In the GO-term landscape, both N- and ade- induced broad stress-response programmes. However, purine-starved cells showed enrichment of trehalose-related processes, whereas nitrogen starvation showed downregulation of the pentose-phosphate pathway (Figures 8D and 5B). These differences are relevant because survival after desiccation requires both protection during drying and recovery after rehydration, including repair of oxidised proteins, membrane restoration, and re-establishment of redox balance. Relative preservation of pentose-phosphate pathway transcription could support NADPH-dependent redox recovery, while trehalose-related transcription and trehalose accumulation could support proteostasis during drying and rehydration. Thus, ribosome biogenesis and rRNA processing repression form a shared starvation backbone in both N- and ade- conditions (Figure 8C), but this shared translation-related repression is not sufficient to explain the desiccation phenotype. The stronger desiccation tolerance of purine-starved cells likely depends on additional metabolic and stress-protection modules that distinguish desiccation-ready states from more general starvation-induced stress resistance.

### Possible signalling pathways behind the purine-starvation response

Our results indicate that, when facing a purine shortage, budding yeast appears to coordinate several parts of its starvation response. Although our data suggest a link to the TOR signalling pathway, the precise signalling route that connects purine depletion to this starvation-like response remains unknown. One possibility is that changes in adenine nucleotide pools contribute to the signal, because purine starvation is expected to affect ATP and GTP metabolism, nucleic acid synthesis, and cofactor metabolism. Consistent with this, purine-starved cells showed transcriptional changes in genes linked to energy metabolism and mitochondrial function. Several highly upregulated transcripts in ade- conditions, including *GLD3, FMP16*, and *OM45*, are consistent with changes in redox metabolism and mitochondrial organisation (Appendix 3). For example, *OM45* and *FMP16* point to altered mitochondrial organisation or respiratory adaptation, whereas *GLD3* may reflect metabolic and redox remodelling.

Purine limitation may also affect DNA replication and genome stability, because purines are required for DNA synthesis. Todeschini et al. (2005) reported that severe adenine limitation increases Ty1 retrotransposon transcripts. We did not detect a clear Ty1-transcription response in our transcriptome dataset, but our data do not exclude changes in Ty1 activity under ade- cultivation conditions. Together, the repression of ribosome biogenesis, changes in carbon and mitochondrial metabolism, G1/0 accumulation, and increased stress resistance point to involvement of nutrient-signalling pathways. TOR and PKA pathways are plausible candidates because they regulate translation, carbon metabolism, mitochondrial function, cell-cycle progression, and stress resistance. This interpretation fits well with the evidence from mammalian cells that reduced purine nucleotide availability can downregulate mTORC1 signalling, even though the relevant yeast mechanism may differ (Hoxhaj et al., 2017). However, previous work showed that rapamycin did not further change the stress resistance caused by purine starvation in *ade8Δ* cells (Kokina et al., 2021), indicating that the relationship between TORC1 inhibition and the purine-starvation phenotype is not straightforward. Consistent with this complexity, deletion of individual candidate transcription factors did not uniformly abolish the purine-starvation-associated recovery phenotype (Figure 9). The effects were strongly background- and medium-dependent, with *gcn2Δade4Δ* under ade- and *gln3Δade4Δ* under N- showing particularly delayed recovery. Thus, the response is unlikely to depend on a single tested regulator, but instead appears to involve a wider nutrient-signalling network. The exact signalling route, therefore, remains unresolved.

Since purine metabolism is highly conserved in eukaryotes, yeast can help identify cellular responses that may also matter in systems dependent on purine uptake or rapid nucleotide synthesis, such as intracellular parasites (Daignan-Fornier & Pinson, 2019; Kokina et al., 2019). Thus, purine-auxotrophic yeast provides a useful model for defining how eukaryotic cells sense and adapt to purine shortage, especially in systems where purine acquisition or nucleotide synthesis represents a metabolic vulnerability.

## Statement on AI-assisted language editing

During manuscript preparation, the authors used OpenAI ChatGPT, GPT-5.5 Thinking model for language editing and stylistic polishing of author-written text. The tool was not used to generate original research data, perform analyses, create figures, interpret results independently, or produce references. All scientific content, interpretations, and final wording were reviewed and approved by the authors, who take full responsibility for the manuscript.

## Acknowledgements

The study was supported by the Latvian Science Council Project LZP-2021/1-0522 “From purine depletion to resilient phenotype – dissecting the mechanisms”

Zane Ozoliņa was supported by the ERDF Project No. 1.1.1.8/1/24/I/003 “Strengthening the Research and Development Capacity of Doctoral Studies at the University of Latvia in the Fields of Smart Specialisation.”

