## Supplementary figures and images for "Position of *de novo* purine biosynthesis gene disruptions shapes purine-starvation phenotypes in *Saccharomyces cerevisiae*"

### Appendix 3

#### Slide 1
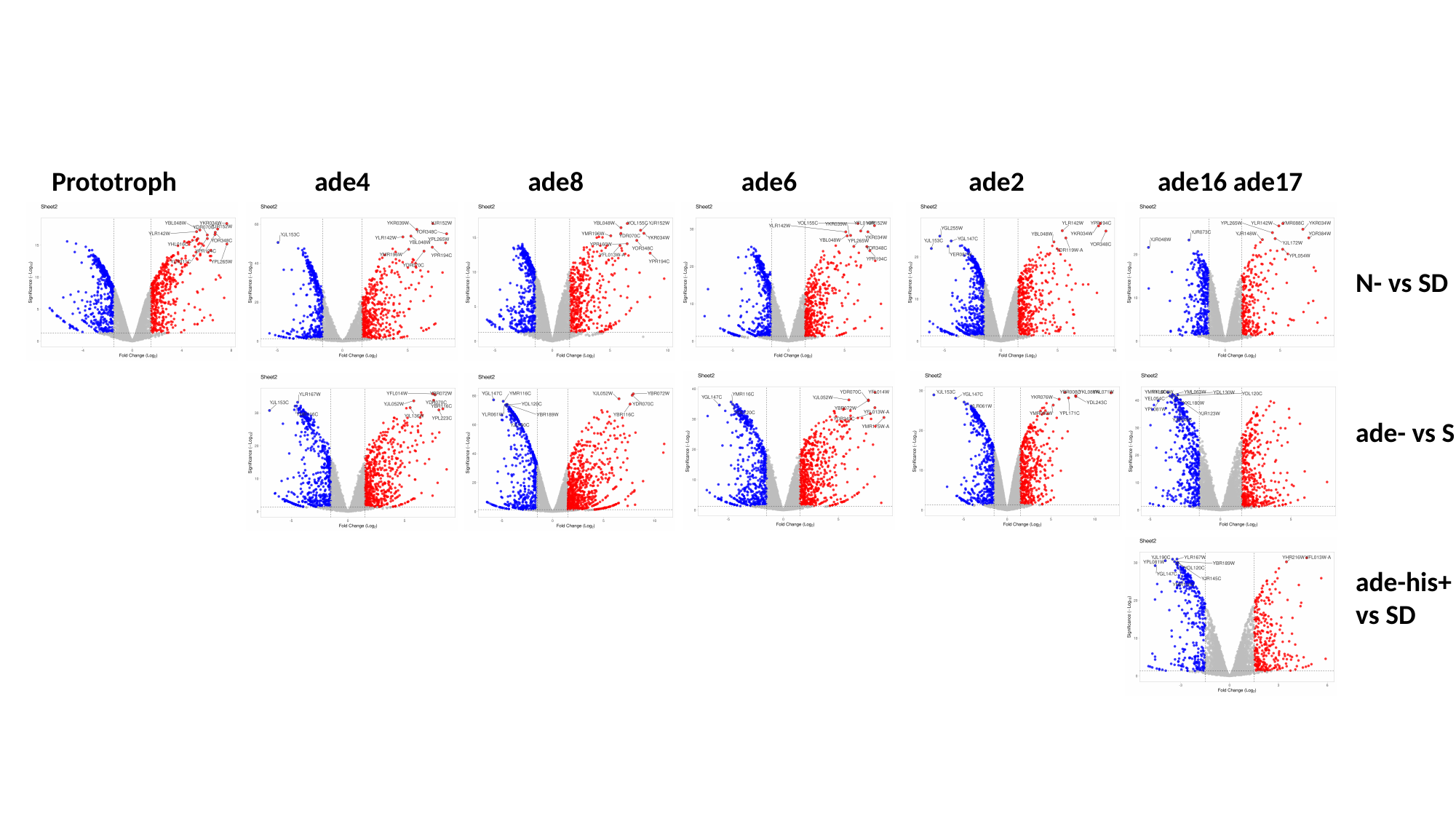

Prototroph
ade4
ade8
ade6
ade2
ade16 ade17
N- vs SD
ade- vs SD
ade-his+
vs SD
